# Loss of BCL-3 sensitises colorectal cancer cells to DNA damage, revealing a role for BCL-3 in double strand break repair by homologous recombination

**DOI:** 10.1101/2021.08.03.454995

**Authors:** Christopher Parker, Adam C Chambers, Dustin Flanagan, Tracey J Collard, Greg Ngo, Duncan M Baird, Penny Timms, Rhys G Morgan, Owen Sansom, Ann C Williams

## Abstract

**Objective:** The proto-oncogene BCL-3 is upregulated in a subset of colorectal cancers (CRC) and increased expression of the gene correlates with poor patient prognosis. The aim is to investigate whether inhibiting BCL-3 can increase the response to DNA damage in CRC.

**Design:** The function of BCL-3 in DNA damage response was studied *in vitro* using siRNA and CRISPR-Cas9 genome editing and *in vivo* using *Bcl3*^*-/-*^ mice. DNA damage induced by γ-irradiation and/or cisplatin was quantified using H2AX and RAD51 foci, repair pathways investigated using HR/NHEJ assays and treatment with the PARP inhibitor olaparib.

**Result:** Suppression of BCL-3 increases double strand break number and decreases homologous recombination in CRC cells, supported by reduced RAD51 foci number and increased sensitivity to PARP inhibition. Importantly, a similar phenotype is seen in *Bcl3*^*-/*-^ mice, where the intestinal crypts of these mice exhibit sensitivity to DNA damage and a greater number of double strand breaks compared to wild type mice. Furthermore *Apc*.*Kras*mutant x *Bcl3*^*-/*-^ mice exhibit increased DNA damage and reduced RAD51+ cells compared to their wild type counterparts when treated with cisplatin.

**Conclusion:** This work identifies BCL-3 as a regulator of the cellular response to DNA damage and suggests that elevated BCL-3 expression could increase resistance of tumour cells to DNA damaging agents including radiotherapy. These findings offer a rationale for targeting BCL-3 in CRC as an adjuvant to conventional therapies and suggest that BCL-3 expression in tumours could be a useful biomarker in stratification of rectal cancer patients for neo-adjuvant chemoradiotherapy.

## Introduction

Rectal cancers constitute around 30% of colorectal malignancies (CRUK). Treatment of rectal cancers differs to that of colonic cancer. Whilst early rectal cancers can be treated with local surgical excision, advanced disease requires surgical resection often with the addition of neoadjuvant therapy to achieve cure. Neo-adjuvant therapy in locally advanced rectal cancer consists of either radiotherapy (short-course radiation), chemoradiotherapy (long-course chemoradiotherapy, LCCRT) or more recently total neoadjuvant therapy (TNT) such as short-course radiotherapy and systemic chemotherapy, as studied in the Rapido trial (1), the aim being to reduce the risk of local or systemic recurrence in these high risk tumours. Importantly, the tumour response to LCCRT is variable (2-7); understanding this variation is critical to improving outcomes in locally advanced rectal cancer.

The ionizing radiation (IR) used in radiotherapy causes a multitude of DNA lesions which result in its anti-tumour effect. Of these lesions, the double strand break (DSB), where the phosphodiester backbone of both strands of DNA are broken in close proximity, is the most cytotoxic and thought to be the primary mode by which radiotherapy reduces tumour growth and survival (8). Following IR, DNA damage repair (DDR) pathways are commonly aberrantly activated in cancer cells leading to tumour cell survival. There are two major DDR pathways by which DSBs can be repaired, non-homologous end joining (NHEJ) and homologous recombination (HR). NHEJ involves recognition of damage by KU proteins which activate DNA-PKcs, followed by repair of the break by end processing enzymes, DNA polymerases and DNA ligase IV (9). This process is fast and active throughout the cell cycle but is error prone. In contrast HR retains DNA sequence fidelity, using a sister chromatid as a template for repair, restricting the process to S and G2 phases of the cell cycle. In this process, the end is processed by the MRN complex (10) CtIP (11), EXO1 and DNA2(12), where the MRN complex also acts as a scaffold for activation of ATM, the major kinase involved in double strand break signalling (13). This processing produces single stranded DNA, initially bound by RPA which is subsequently displaced in the formation of a RAD51 nucleoprotein filament mediated by BRCA2 (14). This filament can then invade homologous DNA which is used as a template for DNA synthesis to restore the damaged sequence.

BCL-3 is a proto-oncogene, highly expressed in a number of solid tumours (15) and is designated as an atypical member of the IκB family owing to its ability to activate or repress transcription of p50 and p52 NF-κB subunits (16). Interactions between BCL-3 and co-regulatory proteins such as TIP60, JAB1, BARD1, PIR, HSP70, HDACs, CTBP1 and 2 and β-catenin (17-20) have been observed, offering additional modes of transcriptional regulation by BCL-3. BCL-3 is implicated in many of the ‘hallmarks of cancer’ including sustaining proliferative signalling, activation of invasion and metastasis and evasion of apoptosis (reviewed in (16)). In CRC, BCL-3 was shown to promote cancer cell growth and survival by phosphoinositide 3-kinase (PI3K) and mammalian target of rapamycin (MTOR) mediated activation of the AKT pathway (21). In addition, it can promote colorectal cancer cell proliferation through regulation of Cyclin D1 (22) and stabilisation of c-MYC (23). More recent work has demonstrated a further role for BCL-3 in cancer, promoting the stem cell phenotype in CRC, with implications for therapeutic resistance (20). Clinically, BCL-3 expression is negatively correlated with survival in CRC patients, where strong nuclear staining of BCL-3 observed in tissue microarrays was associated with reduced patient survival (24) and additionally shown to be independent of tumour stage (20).

Given the poor outcome of patients with high BCL-3 expressing tumours and its previously characterised function in enhancing tumour cell survival, we hypothesised that BCL-3 expression could additionally result in therapeutic resistance of tumours. Here we present data identifying a novel role for BCL-3, where inhibiting BCL-3 expression sensitizes CRC cells to irradiation-induced cell death *in vitro* by reducing HR and show that Bcl-3^-/-^ mice are sensitized to DNA damage inducing agents *in vivo*. This work furthers our understanding of therapeutic resistance in CRC and offers a rationale for targeting BCL-3 as an adjuvant to conventional therapies, particularly in the setting of neoadjuvant therapy for locally advanced rectal cancer.

## Methods

### Cell lines and cell culture

The human rectal carcinoma derived cell line SW1463, and colon carcinoma derived cell lines LS174T and LoVo were obtained from the American Type Culture Collection (ATCC), HCA7 (also colon cancer derived) cells were a kind gift from Sue Kirkham, Imperial College London. The U2OS human osteosarcoma derived cells were a kind gift from Dr Anna Chambers, University of Bristol. All cell lines were cultured in Dulbecco’s Modified Eagle’s Medium (DMEM, Sigma Aldrich, UK) supplemented with 10% foetal bovine serum (Sigma Aldrich, UK), 2mM glutamine, 100 U/mL penicillin and 100 µg/mL streptomycin (Invitrogen). Cells were maintained at 37° C in a dry incubator in 5% CO_2_.

### Gene knockdown with RNAi

Cells were transiently transfected with small interfering RNA (siRNA, 50nM, Dharmacon, UK) using Lipofectamine RNAi MAX (Thermo Fisher Scientific, Paisley, UK). BCL-3 was knocked down using a single siRNA sequence, confirmed with a second sequence (data not shown).

### Generation of BCL3 knockout cells with CRISPR-Cas9 ^D10A^ genome editing

CRISPR-Cas9 ^D10A^ based genome editing was performed according to the method developed by Chiang et al. (25) using the All-In-One vector, provided as a kind gift by Dr Paul Bishop, University of Bristol. Knockout of BCL-3 was confirmed by Sanger sequencing of edit site and absence of BCL-3 protein in western blot.

### Irradiation

Cells were seeded 48 hours prior to irradiation. Irradiation was performed with a Cs137 source in a RX30/55M Irradiator (Gravatrom Industries Ltd.) For RAD51 foci assays, cells were treated with 3µg/mL aphidicolin (Sigma-Aldrich, UK) immediately prior to irradiation to prevent progression from S-phase to G2 phase.

### Crystal Violet Cell Viability Assay

Cells were seeded into and treated in 6 well plates and grown for 48 hrs before treatment. At 14 days after seeding, cells were fixed in 4% paraformaldehyde for 10 minutes and stained with 0.5% w/v crystal violet solution (Sigma-Aldrich, UK) for 10 minutes. Bound crystal violet was eluted with 500μL 10% acetic acid and absorbance measured at 595nm.

### Immunofluorescence

Cells grown on glass coverslips were fixed with 4% paraformaldehyde for 10 minutes and permeabilised with 0.1% Triton-X 100. To prevent non-specific antibody binding, slides were blocked with 1% BSA. Cells were incubated with primary antibody at room temperature for one hour (phospho-histone H2A.X (1:10000, Millipore #05-636); RAD51(1:1000, Abcam); CENPF (1:500, Abcam #5) diluted in 1% BSA. After incubation with secondary antibody (Molecular Probes, Invitrogen, UK) the nuclei were stained 100ng/µL DAPI. Coverslips were mounted with Mowiol 4-88 (Sigma-Aldrich, UK). Cells were imaged with a Leica DMI6000 inverted epifluorescence microscope with 100x lens and Photometrics Prime 95B sCMOS camera or a Leica SPE single channel confocal laser scanning microscope with 40x lens and Leica DFC365FX monochrome digital camera.

### Immunoblotting

Whole cell lysates were made and subjected to SDS-PAGE/western blotting as previously described (26) using the following antibodies: BCL3 (1:2000, Proteintech #, 23959,); CHK2 (1:1000, Millipore #05-649); phospho-CHK2 (1:1000, Cell Signaling Technology, #160); α Tubulin (1:10000, Sigma Aldrich, #T9026).

### Clonogenic Assay

1×10^4^ HCA7 cells were seeded per T25 flask and Olaparib (AZD2281, Selleckchem, UK) was added at indicated concentrations. After 12 days growth, cells were fixed with 10% neutral buffered formalin and stained with 1% Methylene blue (Sigma-Aldrich, UK) for counting number of colonies.

### I-SceI DNA repair assay

On day one, U2OS reporter cell lines (kind gift from Prof J Stark, USA (27)) were transfected with siRNAs (40nM) using lipofectamine RNAiMAX (Thermo Fisher Scientific) and transferred into new plates on day two. Cells were transfected on day three using Dharmafect kb (Dharmacon, UK) with a plasmid pCBASceI to induce I-SceI or with a transfection control plasmid mCherry2-C1. In addition, siRNAs (20nM) were co-transfected to enhance knockdown of the target protein. Cells were analysed on day five using a LSRFortessa™ cell analyser (BD Biosciences, UK). The percentage of GFP positive cells were normalised with percentage of m-cherry positive cells to obtain the DNA repair efficiencies.

### Mouse studies

All animal experiments were performed in accordance with UK Home Office regulations (under project licence 70/8646) and adherence to the ARRIVE guidelines and were subject to review by the Animal Welfare and Ethical Review Board of the University of Glasgow. All mice were maintained on a mixed C57BL/6 background. Mice were housed in conventional cages in an animal room at constant temperature (19–23 °C) and humidity (55%±10%) under a 12-h light–dark cycle and were allowed access to standard diet and water ad libitum. Mice of both genders, aged 2-6 months, were induced with a single intraperitoneal (i.p) injection of 2 mg tamoxifen (#T5648 Sigma-Aldrich,UK) as indicated. The transgenes/alleles used for this study were as follows: *Bcl3*^*-/-*^ (28), *VilCre*^ER^ (29) *Apc*^fl^ (30) and *Kras*^G12D^ (31). For regeneration studies mice were given whole-body irradiation with γ-rays (10 Gy). For treatments, mice were dosed using the following regimens: Cisplatin (5 mg/kg, single intraperitoneal injection), Olaparib (50 mg/kg, once daily oral gavage) and Wee1 inhibitor ((Wee1i) 90 mg/kg, daily oral gavage). All of the listed drugs were made up in 0.5% Hydroxypropyl Methylcellulose (HPMC) and 0.1% Tween-80.

### RNA *in situ* hybridisation

*In situ* hybridisation for *Lgr5* mRNA ((#311838, Advanced Cell Diagnostics) was performed using RNAscope 2.5 LS Reagent Kit–BROWN (Advanced Cell Diagnostics) on a BOND RX autostainer (Leica) according to the manufacturer’s instructions.

### Mouse immunohistochemistry

Intestines were flushed with water, cut open longitudinally, pinned out onto silicone plates and fixed in 10% neutral buffered formalin overnight at 4°C. Fixed tissue was rolled from proximal to distal end into swiss-rolls and processed for paraffin embedding. Tissue blocks were cut into 5 μm sections and stained with haematoxylin and eosin (H&E). Immunohistochemistry (IHC) was performed on formalin-fixed intestinal sections according to standard staining protocols. Primary antibodies used for IHC were against: BrdU (1:150, BD Biosciences, #347580), Rad51 (1:100, Abcam, #133534), Cleaved caspase 3 (1:500, Cell Signaling Technology, #9661) and H2ax (1:50, Cell Signaling Technology, #9718). Representative images are shown for each staining.

## Results

### BCL-3 depletion sensitises CRC cells to gamma irradiation

To determine whether targeting BCL-3 expression could increase the sensitivity of colorectal cell lines to gamma irradiation, four high BCL-3 expressing CRC cell lines (colon HCA7, LS174T, LoVo and rectal SW1463,) were transiently transfected with BCL-3 siRNA, seeded in a 6 well plate and irradiated 48 hrs later with gamma irradiation (1 and 2.5 Gy). After 14 days, viable cells were measured by crystal violet staining (Figure 1, A-D). This revealed that in HCA7 and LS174T cells, BCL-3 knockdown caused lower viability at both 1 and 2.5 Gy whilst in the rectal SW1463 cells, viability was only significantly reduced at the lower dose. However, in LoVo cells, there was no significant decrease in viability when BCL-3 was depleted. These data suggest that, at least in a subset of CRCs, the presence of BCL-3 could be promoting radio-resistance enhancing tumour cell survival.

**Figure 1.**
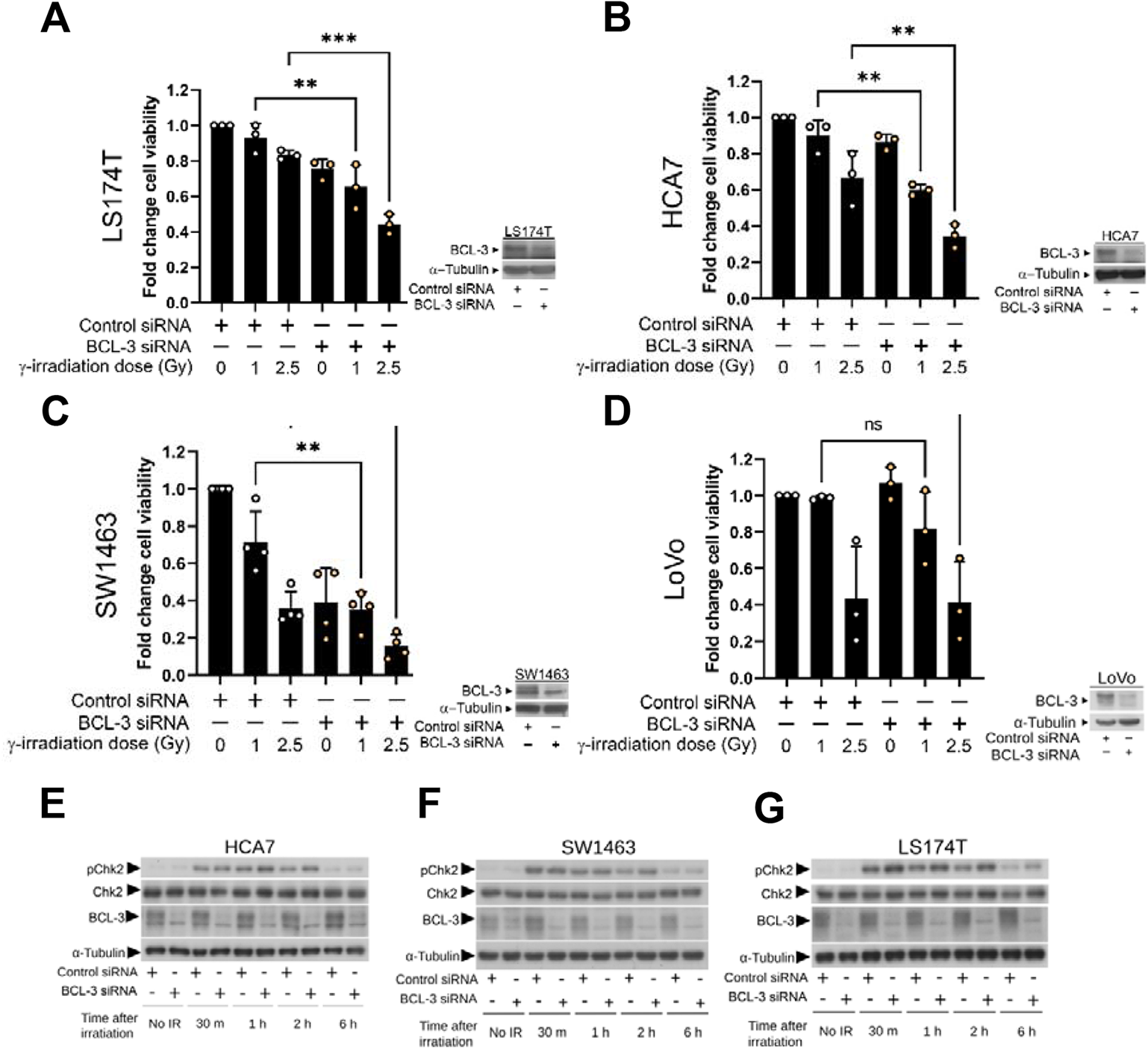
BCL-3 depletion increases sensitivity of colorectal cancer cell lines to gamma irradiation. Four colorectal cancer cell lines, LS174T (**A**), HCA7 (**B**), SW1463 (**C**) and LoVo (**D**) were treated with BCL-3 or control siRNA. 48 hours later, cells were treated with indicated dose of gamma radiation. At 14 days after seeding, viable cells were stained with crystal violet and quantified by absorbance. Knockdown was confirmed by western blot. Data from three independent experiments, statistical significance determined using ANOVA. BCL-3 depletion increases ionising radiation induced double strand break number and downstream signalling. HCA7 (**E**), SW1463 (**F**) and LS174T (**G**) cells were treated with BCL-3 or control siRNA and given a 2.5Gy dose of ionising radiation 48 hours later. Cell lysates were taken at time points indicated and subjected to western blotting for total and phosphorylated CHK2. Tubulin was used as a loading control. Data representative of three experimental repeats.

### Loss of BCL-3 increases DSB associated signalling and DSB number after γ- irradiation in sensitive human colorectal cells

Because DSB generation is the primary mode of cell death/quiescence in irradiated cells, we hypothesised that suppression of BCL-3 expression reduced the capacity of the cell to repair DSBs. ATM is the primary transducer of signalling that communicates formation of DSBs from the proteins that recognise the break to effectors of repair (32). After DNA damage, ATM is rapidly activated by autophosphorylation (33) which initiates the phosphorylation and activation of a multitude of proteins involved in DNA repair and cell cycle arrest including H2AX and CHK2 (34-36). To determine whether repression of BCL-3 changed signalling downstream of DSB formation, the sensitive CRC cells (LS174T, HCA7 and SW1464) were again transiently transfected with BCL-3 siRNA and treated with IR (2.5Gy) 48 hours later. Lysate samples were then taken between 0-6 hours post irradiation. Western blotting was used to measure activation of CHK2 using phosphorylation specific antibodies (Figure 1, E-G). In all three cell lines there was rapid induction of CHK2 phosphorylation after irradiation (within 30 minutes). When BCL-3 was suppressed in the cells, phosphorylation of CHK2 was further increased at one and two hours after irradiation in all three cell lines, suggesting more damage had occurred following BCL-3 suppression. To investigate this further, foci of γ-H2AX, a marker that directly corresponds to DSB number, were quantified in HCA7 cells (chosen for amenability to imaging). BCL-3 was knocked down in HCA7 cell by transient siRNA transfection and cells were irradiated with 2.5 Gy 48 hours later. Cells were fixed at points 0-6 hours after irradiation and stained with antibody for visualisation of γ-H2AX. Interestingly, BCL-3 depleted HCA7 cells exhibited significant increased γ-H2AX foci number in the unirradiated cells (Figure 2A), suggesting more DSB are present in the cells in which BCL-3 is suppressed. On induction of DSB using 2.5Gy γ-irradiation, this difference was further increased, with the BCL-3 siRNA transfected cells having significantly higher levels of γ-H2AX foci for up to two hours after irradiation. This increase in DSB number suggests that depletion of BCL-3 increases spontaneous break formation and/or reduces rate of DSB repair. By six hours after irradiation the difference in γ-H2AX foci number is lost, with the number of γ-H2AX foci in BCL-3 siRNA being similar to that of the siRNA control cells at the two-hour time point. This effect was confirmed in the rectal cancer SW1463 cells 30 min after irradiation as well (Figure 2B).

**Figure 2.**
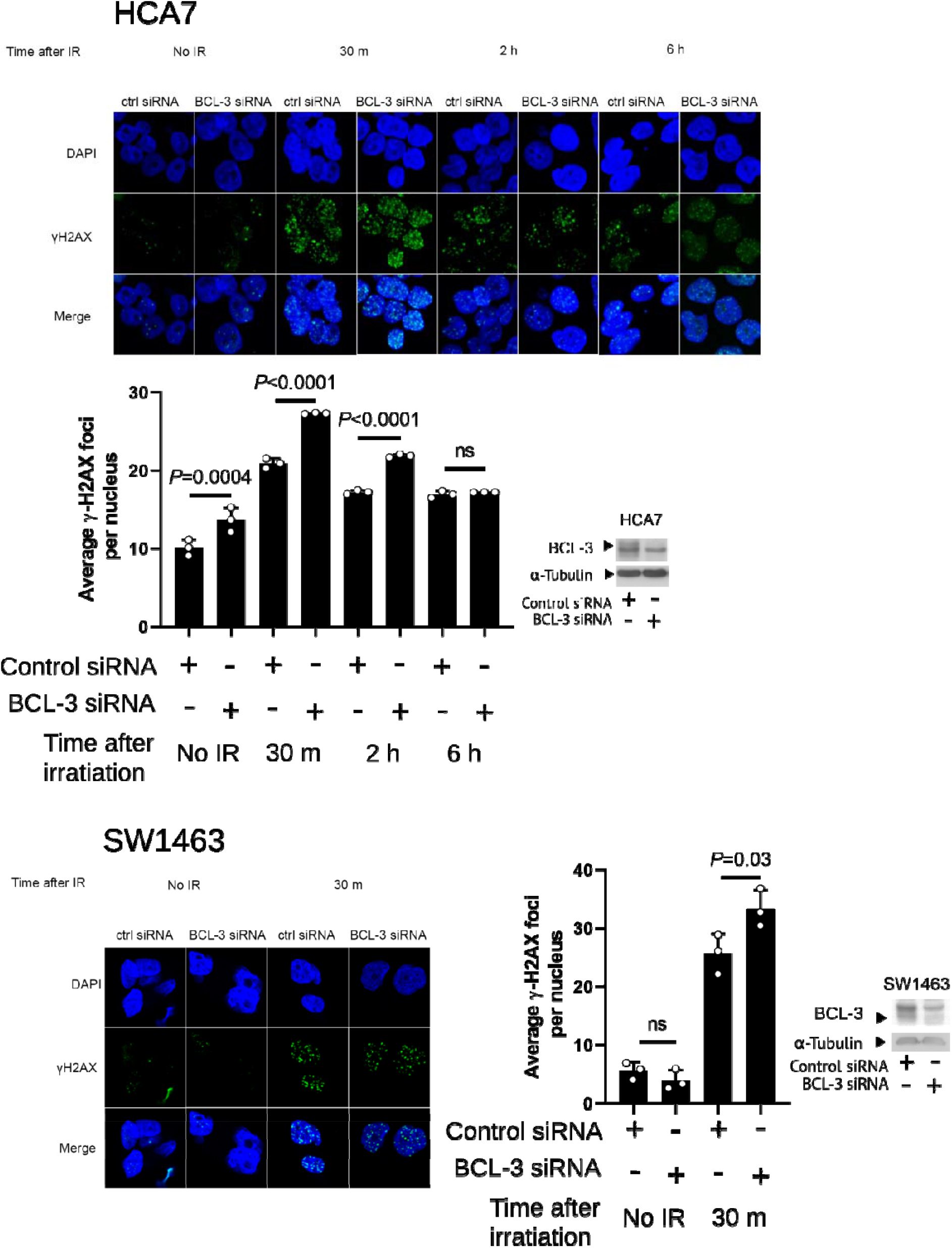
BCL-3 depletion increases ionising radiation induced double strand break number. HCA7 (**A**) and SW1463 (**B**) cells were treated with BCL-3 siRNA and negative siRNA and given a 2.5Gy dose of ionising radiation 48 hours later. Cells were fixed at time points indicated after irradiation and stained for γ-H2AX. Data from three independent experiments analysing > 30 nuclei per repeat. statistical significance determined using ANOVA.

### Phosphoproteomics reveal a role for BCL-3 in the DNA damage response

Having observed an increase in DSBs in cells with reduced BCL-3 expression, we sought to investigate the possible mechanism. To take an unbiased view of potential pathways changed following loss of BCL-3, TMT-phosphoproteomics was performed on whole cell HCA7 lysates. For these experiments BCL-3 was deleted by CRISPR-Cas9^D10A^ gene editing (25), resulting in production of a BCL-3 knockout clone (KO) and a pool of control cells (NKO) which had undergone the knockout protocol but in which BCL-3 was not deleted (see supplementary Figure 1). Wildtype (WT), NKO and KO HCA7 cells were seeded at equal density and lysates were taken after 48 hours of growth. Lysates were trypsin digested and the resulting peptides differentially labelled prior to MS/MS analysis. The data produced were subject to a Sequest search against the Uniprot human database and filtered at a 5% false discovery rate. Protein phosphorylation that was affected by BCL-3 deletion was identified as that with significantly different (p<0.05) abundance between the NKO and BCL-3 KO populations after excluding those proteins that also changed between WT and NKO cells. This showed an increased abundance of 258 phosphopeptides and decreased abundance of 146 phosphopeptides in the BCL-3 KO cells relative to the NKO cells. Ingenuity Pathway Analysis (IPA, QIAGEN) revealed that a number of pathways related to DNA damage repair (DDR) and control were changed by the loss of BCL-3 (Figure 3A). Of interest, changes in “the role of BRCA1 in DNA Damage Response” canonical pathway suggest that phosphorylation of proteins directly involved in DNA damage repair could be changed in BCL-3 KO cells. This includes DNA repair proteins RAD50, RFC1 and FANCD2. Examining changes in biological functions by IPA showed a predicted increase in DNA damage in the BCL-3 KO cells relative to the NKO cells. This analysis was repeated in another BCL-3 KO colorectal cell line, demonstrating similar changes in pathways involved in homologous recombination (Supplementary Figure 2).

**Figure 3.**
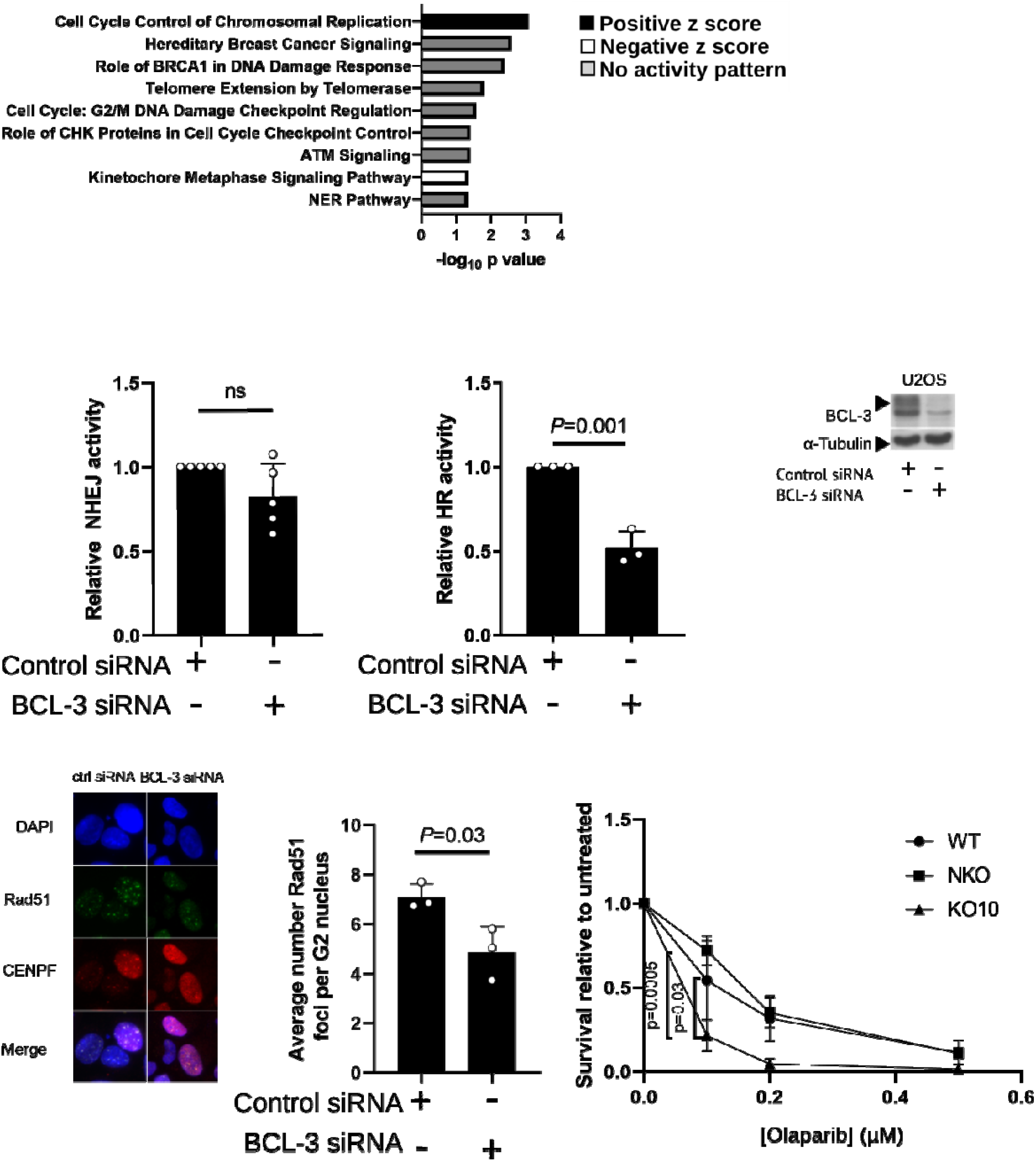
BCL-3 knockdown reduces rate of homologous recombination in U2OS and colorectal cancer cells. (**A)** Canonical pathway enrichment analysis with of differentially expressed phosphoproteins when comparing *BCL-3*^-/-^ vs CRISPR control HCA7 cells. Level of enrichment is shown by -log(p-value). Z-score indicates the of predicted activation state of the canonical pathway. Blue indicates a negative z-score and pathway inhibition. Orange indicates a positive z-score and pathway activation. BCL-3 was knocked down in EJ5-GFP (**B**) and DR-GFP (**C**) U2OS cells transfected with a I-SceI expression plasmid. NHEJ (B) and HR (C) activity were measured by percentage of GFP positive cells in FACS, expressed relative to control siRNA population. Significance measured with two tailed T test. **(D)** Representative western blot of BCL-3 knockdown by siRNA in U2OS cells in B,C and E. (**E**) RAD51 foci were counted in CENPF positive cells two hours after 2.5 Gy treatment in U2OS cells treated with indicated siRNA. Significance measured by two tailed T-test. (**F**) WT, CRISPR control (NKO) and BCL-3 knock out (KO) cells were seeded at equal density and treated six hours later with olaparib. Number of colonies formed was counted two weeks after seeding, represented as relative mean survival, error bars represent 1SD. Data from three independent experimental repeats, significance measured by one way ANOVA.

### BCL-3 promotes DSB repair by Homologous Recombination

To investigate whether loss of BCL-3 expression regulated DSB repair in cells, we used EJ5-GFP and DR-GFP assays to measure the effects of BCL-3 depletion on rates of NHEJ and HR respectively (27). These assays use U2OS cell lines with an integrated cassette for GFP, the sequence of which is interrupted by a restriction site for the restriction enzyme I-SceI. Expression of I-SceI in these cells generates double strand breaks which can be repaired only by the pathway specific to the cell line (HR; DR-GFP or NHEJ; EJ5-GFP), generating GFP+ cells that are subsequently quantified by flow cytometry. BCL-3 was depleted in the U2OS EJ5-GFP cells by siRNA transfection, followed 48 hours later by I-SceI transfection. Flow cytometry subsequently revealed that in there was no significant difference in reporter activity (Figure 3B) suggesting that changes in BCL-3 expression did not affect NHEJ. However, there was a marked and significant decrease in reporter activity in DR-GFP cells when BCL-3 was supressed (Figure 3C), consistent with drop in HR and in concordance with HR being the most changed pathway in the phosphoproteome.

RAD51 is a HR specific repair factor that accumulates at repair foci in S and G2 phase cells. To further investigate whether suppression of BCL-3 reduced HR as shown by the DR-GFP reporter assay, quantitation of RAD51 foci by immunofluorescence in U2OS cells was used as a measure of HR activity (37). This cell line was selected for its suitability in immunofluorescent imaging of RAD51 after unsuccessful attempts to stain Rad51 in HCA7 cells. To restrict the analysis to cells in which HR is active, cells were co-stained with CENP-F, a marker of G2 and S phase cells. BCL-3 was knocked down in U2OS cells by transient siRNA transfection and the cells were irradiated with 2.5 Gy 48 hours later. The cells were fixed two hours after irradiation to capture the peak in foci number (38) and RAD51 foci were visualised with immunofluorescense using a recombinant RAD51 antibody. Foci were counted in a minimum of 60 CENP-F positive nuclei per experimental repeat; the number of RAD51 foci were significantly reduced in BCL-3 depleted U2OS cells (Figure 3E), supporting the role of BCL-3 in HR.

### BCL-3 depletion sensitises CRC Cells to PARP inhibitors

Cells with a defect in any component of HR are sensitive to treatment with PARP inhibitors, as exploited therapeutically in cancers where BRCA1 or BRCA2 are mutated (39). Inhibition of PARP reduces SSB repair resulting in a greater DSB load when SSBs are converted to DSBs in DNA replication (39-41). The effect of BCL-3 on sensitivity to PARP inhibition was assessed by clonogenic assay where WT, NKO and BCL-3 KO HCA7 were seeded into flasks at 1×10^4^ per flask and treated with olaparib. After 14 days, the number of colonies formed was counted. BCL-3 KO CRC cells were more sensitive to the PARP inhibitor olaparib than control cells, showing reduced colony formation after treatment (Figure 3F), concordant with defective HR in the CRC cells.

### BCL-3 protects the intestinal crypt from DNA damage and oncogenic-induced stress

To translate our findings to a more complex system, we examined the sensitivity and response of the intestinal epithelium following ionising radiation and DNA damage in BCL-3 knockout mice (*Bcl3*^-/-^). In this context, control littermates (WT) and *Bcl3*^-/-^ mice were given a dose of whole body γ-irradiation (10 Gy), triggering a well-defined cascade of cell death (6-24hrs) and epithelial regeneration (72hrs) following damage (42). Compared to WT control, the regenerative capacity of *Bcl3*^-/-^ mice was significantly impaired, as measured by the total number of regenerating crypts and BrdU^+^ crypt cells (Figure 4A-C). Given the increased frequency of DSBs following loss of BCL-3 in CRC cells, we sought to determine whether DSBs are also increased in *Bcl3*^-/-^ intestinal epithelia. In line with previous results (Figure 2), *Bcl3*^-/-^ intestinal crypts showed higher levels of γH2AX foci 24hrs post-irradiation compared to control mice (Figure 4D). Importantly, *Bcl3*^-/-^ mice showed increased cleaved-caspase 3^+^ cells at the same time point (24hrs) compared to control (Figure 4E), suggesting that failure to repair DSBs triggers cell death, and subsequent compromised tissue regeneration. Importantly, cells with defective DDR or deficient HR show sensitivity to drugs that induce DNA damage, such as cisplatin (40, 41). We leveraged this, and treated *Bcl3*^-/-^ mice with cisplatin (5mg/kg), olaparib (50mg/kg) or a wee1 inhibitor (90mg/kg) for 5 days to better understand the function of BCL-3 in regulating DNA damage response. Examination of crypts 24hrs following treatments showed that whilst there was no significant difference in number of BrdU^+^ cells between WT and *Bcl3*^-/-^ mice with any treatment (Supplementary Figure 3), there were significantly more γH2AX foci in *Bcl3*^-/-^ mice compared to WT when treated with cisplatin (Figure 4F, G). Similarly, *Bcl3*^-/-^ mice treated with olaparib had increased γH2AX foci compared to WT control (Supplementary Figure 3), consistent with *in vitro* findings (Figure 3E). Interestingly, there was no change to DNA damage (γH2AX foci) or cell death following wee1 inhibition (Supplementary. Figure 3), which is consistent with its non-HR targeted mechanism of action (43). Taken together, the increased damage seen in *Bcl3*^-/-^ mice with cisplatin and in particular, olaparib treatment is consistent with a defect in HR as seen in BCL3-deficient human CRC cells (Figure 3).

**Figure 4.**
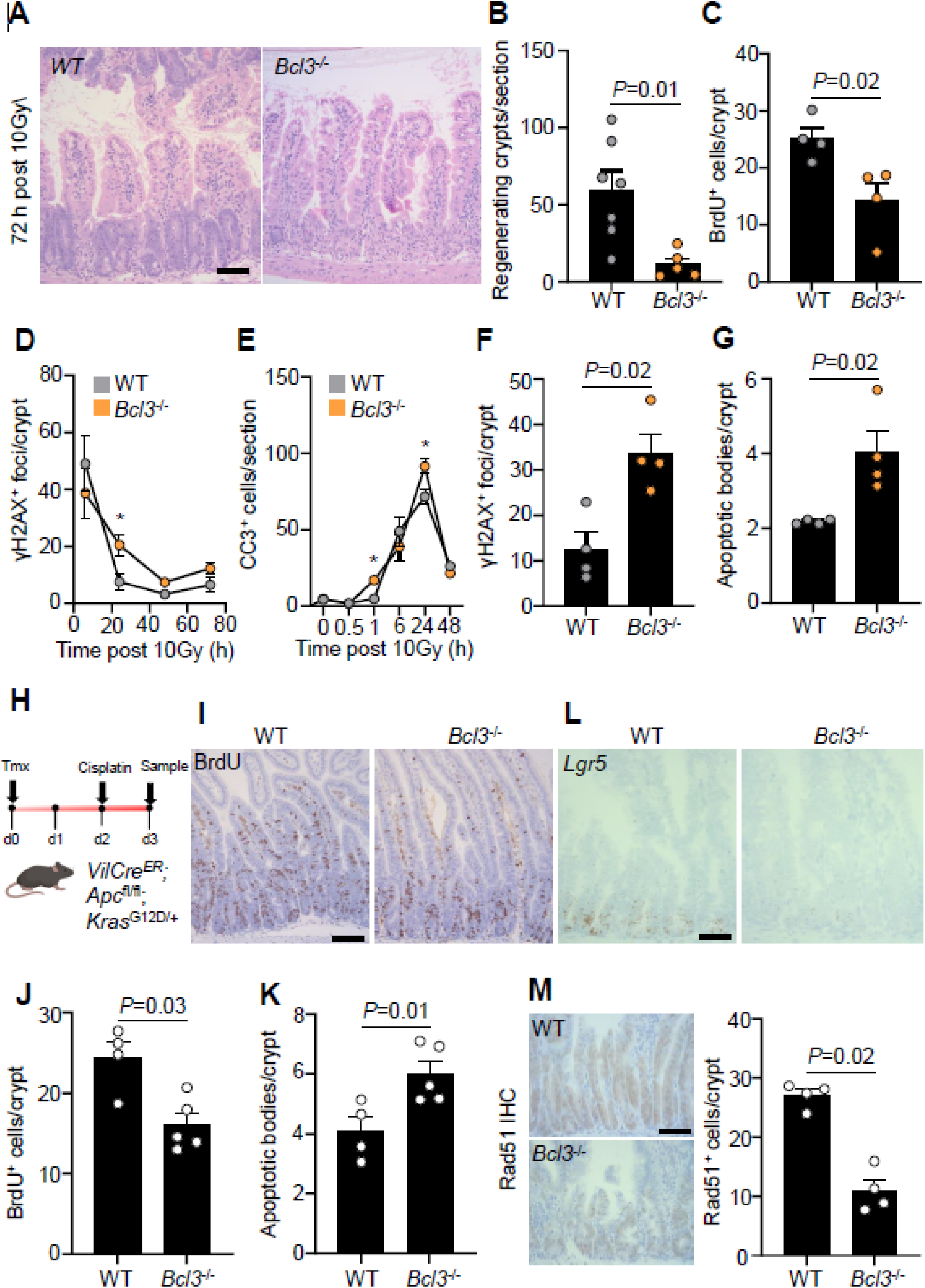
Loss of Bcl3 impairs intestinal regeneration and sensitises the epithelium to DNA damage. (**A**) Representative H&E staining of wild-type (WT) and Bcl3KO (*Bcl3*^-/-^) mice 72 hrs following whole-body irradiation (10 Gy). Scale bar, 100 μm. (**B**) Quantification of regenerating crypts per intestinal cross section from mice described in A. n=7 WT, n=5 *Bcl3*^-/-^. Data are ± s.e.m; Mann–Whitney two-tailed U-test. (**C**) Quantification of BrdU^+^ cells per regenerating crypt from mice described in A. n=4 WT, n=4 *Bcl3*^-/-^. Data are ± s.e.m; Mann– Whitney two-tailed U-test. (**D**) Quantification of γH2AX^+^ foci per crypt over time following whole-body irradiation (10Gy). n=4 WT, n=4 *Bcl3*^-/-^. Data are ± s.e.m; *P=0.01. Mann– Whitney two-tailed U-test. (**E**) Quantification of cleaved caspase-3^+^ (CC3^+^) cells per intestinal cross section over time following whole-body irradiation (10 Gy). n=4 WT, n=4 *Bcl3*^-/-^. Data are ± s.e.m; *P=0.01. Mann–Whitney two-tailed U-test. (**F**) Quantification of γH2AX^+^ foci per crypt 24hrs following single dose of cisplatin (5 mg/kg) in mice of the indicated genotypes. n=4 WT, n=4 *Bcl3*^-/-^. Data are ± s.e.m; Mann–Whitney two-tailed U-test. (**G**) Quantification of apoptotic bodies per crypt in mice described in F. n=4 WT, n=4 *Bcl3*^-/-^. Data are ± s.e.m; Mann–Whitney two-tailed U-test. (**H**) Schematic illustrating tamoxifen induction, treatment regimen and tissue harvest of *VilCre*^*ER*^;*Apc*^fl/fl^;*Kras*^G12D/+^ mice +/- *Bcl3*. (**I**) Representative BrdU staining of tamoxifen-induced *VilCre*^*ER*^;*Apc*^fl/fl^;*Kras*^G12D/+^ (WT) and *VilCre*^*ER*^;*Apc*^fl/fl^;*Kras*^G12D/+^;*Bcl3*^-/-^ (*Bcl3*^-/-^) mice 24 hrs following cisplatin treatment (5 mg/kg). Scale bar, 100 μm. (**J**) Quantification of BrdU^+^ cells per crypt/villus unit in mice described in n=4 WT, n=5 *Bcl3*^-/-^. Data are ± s.e.m; Mann–Whitney two-tailed U-test. (**K)** Quantification of apoptotic bodies per crypt in mice described in I. n=4 WT, n=5 *Bcl3*^-/-^. Data are ± s.e.m; Mann–Whitney two-tailed U-test. (**L**) Representative *Lgr5* (in-situ hybridisation) staining of mice described in I. Scale bar, 100 μm. (**M)** Representative staining (left) and quantification (right) of Rad51^+^ cells in mice described in I. Scale bar, 100 μm. n=4 WT, n=4 *Bcl3*^-/-^. Data are ± s.e.m; Mann–Whitney two-tailed U-test.

To extend and confirm the findings in human CRC cells, we generated mice harbouring common CRC mutations, *Apc* and *Kras*, targeted to the intestinal epithelium (*VilCre*^ER^;*Apc*^fl/fl^;*Kras*^G12D/+^, hereafter WT) and crossed them to *Bcl3*^-/-^ mice (*VilCre*^ER^;*Apc*^fl/fl^;*Kras*^G12D/+^;*Bcl3*^-/-^, hereafter *Bcl3*^-/-^). Both WT and *Bcl3*^-/-^ mice were induced with tamoxifen (2 mg) and treated with a single dose of cisplatin (5 mg/kg) and harvested 24hrs after (Figure 4H). Consistent with cisplatin’s mechanism of action (44), highly-proliferative transit-amplifying cells were significantly reduced in both WT and *Bcl3*^-/-^ mice compared to untreated controls (Figure 4I, Supplementary Figure 3). However, compared to cisplatin-treated WT mice, cisplatin-treated *Bcl3*^-/-^ mice had reduced BrdU^+^ crypt cells with a concomitant increase in apoptosis (Figure 4I-K, Supplementary Figure 3), indicating loss of Bcl3 sensitises *Apc*.*Kras*-mutant cells to chemotherapeutics. We have previously showed Bcl3 promotes stem-cell phenotypes in CRC cells (45), and therefore wanted to determine whether the combination of DNA damage (cisplatin) and Bcl3 deficiency preferentially targets *Lgr5*^+^ intestinal stem cells (46) and/or transit-amplifying progenitor cells. In support of our previous work (20), *Lgr5* expression was markedly reduced in cisplatin-treated *Bcl3*^-/-^ mice, while WT mice retained *Lgr5* expression, suggesting cisplatin and loss of Bcl3 synergise to eliminate both the TA and importantly the Lgr5^+^ ISC populations (Figure 4L). Finally, to investigate whether the increased sensitivity following cisplatin in *Bcl3*^-/-^ mice was due to defective HR, we quantified Rad51^+^ crypt cells following cisplatin. Consistent with previous results (Figure 3D), *Bcl3*^-/-^ mice had reduced Rad51^+^ cells following DNA damage (cisplatin) compared to WT mice (Figure 4M), indicating Bcl3 is required for optimal HR signalling after DNA damage.

## Discussion

Given the relationship between BCL-3 expression levels in CRC and poor patient prognosis (20, 24, 47), we sought to investigate whether BCL-3 plays a role here by altering tumour response to therapeutic agents. Here, we report that loss of BCL-3 sensitises colorectal cells to DNA damaging therapeutic agents both *in vitro* and *in vivo*. Although BCL-3 has been studied in the context of DNA damaging agents such as UV irradiation (48-50) and alkylating agents (51), to our knowledge no investigation have been made of a role for BCL-3 in the DDR. This builds on previous work that has characterised a role for BCL-3 in promoting tumour survival, inhibiting apoptosis through the AKT pathway (21), HDM2 (49, 52-54) and DNA-PK (49). In addition, recent work in our group has also demonstrated BCL-3 promoting the stem cell phenotype in CRC cells, a phenotype associated with therapeutic resistance (20). Here, we observed increased γH2AX foci number upon BCL-3 loss in both CRC cells and mouse epithelium after irradiation, suggesting that this effect is the result of either a greater amount of DNA damage being sustained or that cells with depleted BCL-3 have acquired a defect in the DDR. The observed reduction in HR activity coupled with increased sensitivity to PARP inhibition in BCL-3 depleted cells both *in vitro* and *in vivo* suggests that loss of repair activity is at least in part responsible for greater DSB number.

Several earlier studies could potentially shed light on mechanisms by which BCL-3 may regulate the rate of DNA repair by HR. BCL-3 is thought to bind to a number of proteins involved in DSB repair. Deschend et al. (17) identified interaction of a BCL-3-p50 complex with JAB1, BARD1 and TIP60. BARD1 is the major binding partner of BRCA1 (55) which in turn plays a direct role in HR, directing RAD51 filament formation and inhibiting NHEJ (56). The upregulation of this pathway with loss of BCL-3 seen in phosphoproteomics in BCL-3 KO cells could be to compensate the loss of the interaction between BCL-3 and BARD1. TIP60 is a histone acetyl transferase (HAT) and another key DNA repair protein, required for acetylation and activation of ATM and DNA-PKcs in addition to a role in acetylation of histones H2AX and H4 at DSBs (57). BCL-3 also interacts with the HAT p300/CBP(58) and histone deacetylases (HDACs)(18, 59-61). HDACs are commonly upregulated in CRC and these proteins are required for expression and catalytic activity of a multitude of DNA repair proteins as well as altering chromatin compaction for repair (62). Therefore loss of BCL-3 could be mimicking the radiation sensitivity seen upon knockdown of class I HDACs (63, 64).

The identification of BCL-3 as a HR promoting factor has several implications for treatment of CRC patients. The observed increase in radiosensitivity upon BCL-3 depletion suggests that rectal patients who have higher tumour BCL-3 expression would be less likely to respond to neo-adjuvant radiotherapy, therefore stratification of therapy by BCL-3 expression could avoid unnecessary treatment or necessitate the addition of adjunctive therapies to improve therapeutic response (65). However, we observed that sensitisation to γ-irradiation by BCL-3 knockdown was not universal across the high BCL-3 expressing cell lines tested, a comparison between BCL-3 expression in patient tissue and response to radiotherapy will therefore be required to demonstrate the applicability of BCL-3 as a predictor of therapeutic response.

Adjuvant treatment with novel chemical inhibitors of BCL-3 (66) may allow patients with higher BCL-3 expression to be sensitised to conventional radiotherapy, improving therapeutic response. As demonstrated *in vivo*, this finding would also be relevant to DNA damaging chemotherapy. Several agents that cause DNA damage repaired by HR are used or have been trialled in CRC. As well as platinum based compounds, Irinotecan targets topoisomerase I during DNA replication, causing DSBs (67) whilst Veliparib is a PARP inhibitor and has been used in trials in combination with other drugs (68). Furthermore, there is increasing interest in the concept of BRCAness where factors other than BRCA1 or BRCA2 mutation result in defective HR and an expanding group of proteins are being targeted to increase sensitivity to agents such as PARP inhibitors (69). Our results suggest that targeting BCL-3 during treatment to achieve an HR reduced phenotype would increase the efficacy of these drugs. Alternatively, BCL-3 expression could be used to stratify patients for treatment with these drugs, avoiding their use in more HR proficient cells. Excitingly, given the expression of BCL-3 in a wide range of cancers including breast cancer, these findings could translate into improving therapy response or stratifying treatment in other cancer types beside CRC.

In conclusion we have demonstrated a novel role for BCL-3 in conferring resistance to IR in colorectal cancer cells and in mouse intestine. In addition, we have shown that inhibition of BCL-3 function could increase the therapeutic response to chemotherapeutic drugs as well. We propose that patients with higher BCL-3 expressing colorectal tumours have poorer survival at least in part because it promotes increased HR activity, allowing cells to tolerate and repair DNA damage caused by therapeutic agents.

## Acknowledgements

We thank Prof. Jeremy Stark (City of Hope Comprehensive Cancer Centre, USA) for providing the U2OS EJ5-GFP and DR-GFP reporter cell lines. We thank the Wolfson Bioimaging Facility, University of Bristol for assistance with imaging.

