## Supplemental Figure 1 for "Loss of BCL-3 sensitises colorectal cancer cells to DNA damage, revealing a role for BCL-3 in double strand break repair by homologous recombination"

**A**

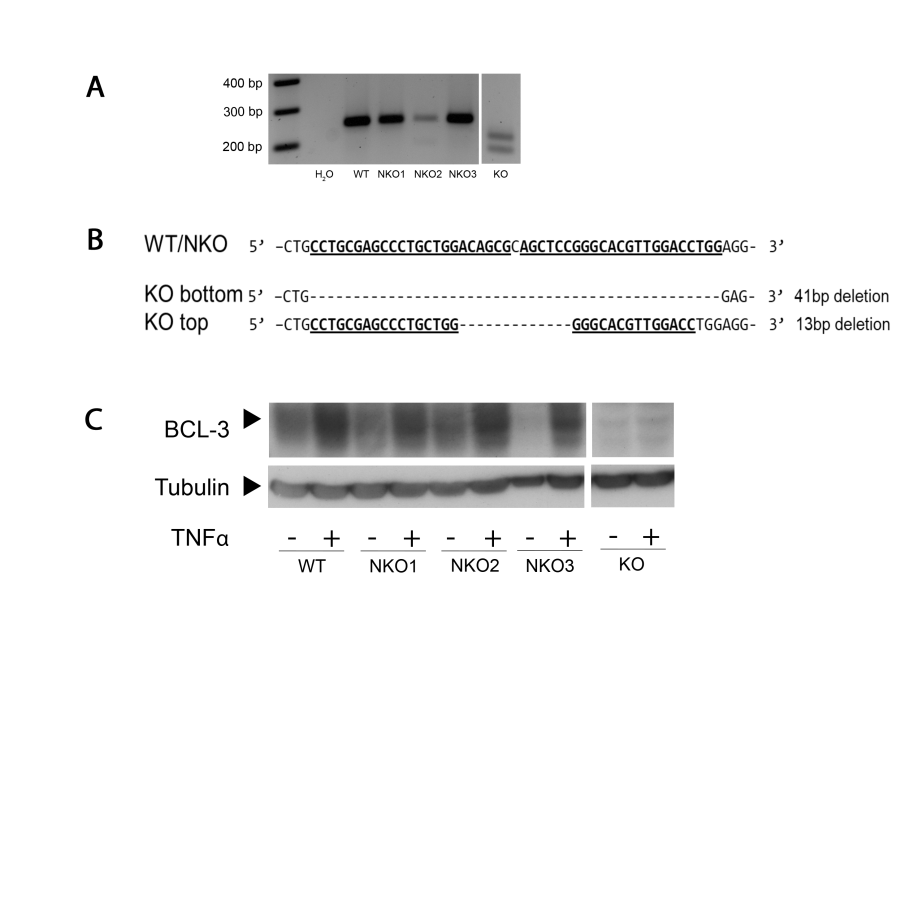

**C**

**B**

**Supplementary Figure 1**. **A**, DNA extracted from HCA7 clones [WT; non- knockout control, (NKO) and knockout, (KO)) after the BCL-3 knockout process underwent PCR with primers complementary to the Cas9^D10A^ target site. PCR products were subsequently separated on a 2% agarose gel by electrophoresis. **B**, PCR products underwent Sanger sequencing demonstrating identical sequences in the WT cells and NKO clones and deletions of 41 base pairs and 13 base pairs in the bottom and top bands of the KO clone. Sequence shown flanks the Cas9^D10A^ target site with sgRNA sequences shown in bold underlined. **C**) BCL-3 expression from WT, NKO and KO clones after TNFα stimulation confirming loss of BCL-3 expression in the KO clone.
