## Supplemental Figure 2 for "Loss of BCL-3 sensitises colorectal cancer cells to DNA damage, revealing a role for BCL-3 in double strand break repair by homologous recombination"

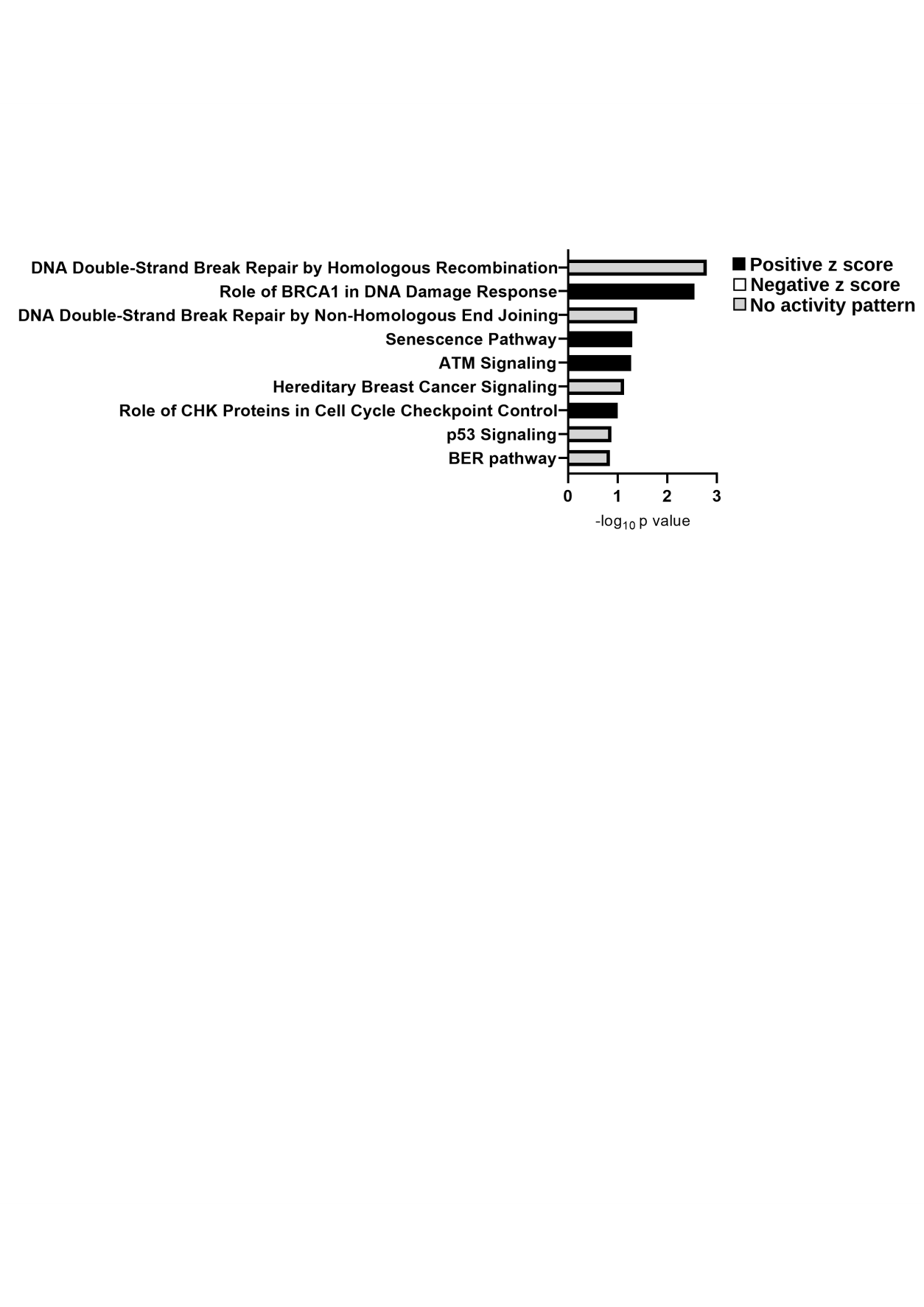

**Supplementary Figure 2**. Canonical pathway enrichment analysis of differentially expressed phosphoproteins when comparing BCL-3 KO vs CRISPR NKO control SW620 cells. Level of enrichment is shown by -log(p-value). Z-score indicates the of predicted activation state of the canonical pathway. Blue indicates a negative z-score and pathway inhibition. Orange indicates a positive z-score and pathway activation.
