## Supplemental Figure 3 for "Loss of BCL-3 sensitises colorectal cancer cells to DNA damage, revealing a role for BCL-3 in double strand break repair by homologous recombination"

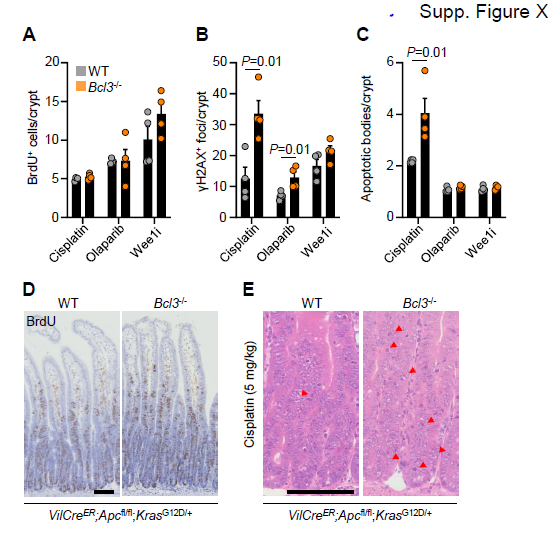
**Supplementary Figure 3. Bcl3 is required to protect the intestinal epithelium against DNA damage.**

**A-C.** Quantification of BrdU^+^ (A)**,** γH2AX^+^ (B) and apoptotic cells (C) per crypt in mice of the indicated genotype 24hrs after the last dose of the indicated treatments (x-axis). n=4 WT, n=4 *Bcl3*^-/-^. Data are ± s.e.m; Mann–Whitney two-tailed U-test. **D**. Representative BrdU staining of *VilCre^ER^;Apc*^fl/fl^*;Kras*^G12D/+^ (WT) and *VilCre^ER^;Apc*^fl/fl^*;Kras*^G12D/+^;*Bcl3*^-/-^ (*Bcl3*^-/-^) mice 3 days post tamoxifen-induction. Scale bar, 100 μm. **E**. Representative H&E staining of tamoxifen-induced *VilCre^ER^;Apc*^fl/fl^*;Kras*^G12D/+^ (WT) and *VilCre^ER^;Apc*^fl/fl^*;Kras*^G12D/+^;*Bcl3*^-/-^ (*Bcl3*^-/-^) mice 24 hrs post cisplatin treatment (5 mg/kg). Red arrowheads indicate apoptotic cells. Scale bar, 100 μm.
